# Environmental biases in the study of ecological networks at the planetary scale

**DOI:** 10.1101/2020.01.27.921429

**Authors:** Timothée Poisot, Gabriel Bergeron, Kevin Cazelles, Tad Dallas, Dominique Gravel, Andrew Macdonald, Benjamin Mercier, Clément Violet, Steve Vissault

## Abstract

Ecological networks are increasingly studied at large spatial scales, expanding their focus from a conceptual tool for community ecology into one that also adresses questions in biogeography and macroecology. This effort is supported by increased access to standardized information on ecological networks, in the form of openly accessible databases. Yet, there has been no systematic evaluation of the fitness for purpose of these data to explore synthesis questions at very large spatial scales. In particular, because the sampling of ecological networks is a difficult task, they are likely to not have a good representation of the diversity of Earth’s bioclimatic conditions, likely to be spatially aggregated, and therefore unlikely to achieve broad representativeness. In this paper, we analyze over 1300 ecological networks in the mangal.io database, and discuss their coverage of biomes, and the geographic areas in which there is a deficit of data on ecological networks. Taken together, our results suggest that while some information about the global structure of ecological networks is available, it remains fragmented over space, with further differences by types of ecological interactions. This causes great concerns both for our ability to transfer knowledge from one region to the next, but also to forecast the structural change in networks under climate change.

## 1 Introduction

Ecological networks are a useful representation of ecological systems in which species or organisms interact (Heleno *et al.* 2014, Poisot & Stouffer *et al.* 2016, Delmas *et al.* 2018), and there has been a recent explosion of interest in their dynamics across large temporal scales (Tylianakis & Morris 2017, Baiser *et al.* 2019), and along environmental gradients (Trøjelsgaard & Olesen 2016, Pellissier *et al.* 2017). As ecosystems are changing rapidly, networks are at risk of undergoing rapid and catastrophic changes to their structure: for example by invasion leading to a collapse (Strong & Leroux 2014, Magrach *et al.* 2017), or by a “rewiring” of interactions among existing species (Bartley *et al.* 2019, Guiden *et al.* 2019, Hui & Richardson 2019). Simulation studies suggest that knowing the structure of the extant network, *i.e.* being able to map all interactions between species, is not sufficient (Thompson & Gonzalez 2017) to predict the effects of external changes, and that data on the species, the local climate and its future projection, are also required.

This change in scope, from describing ecological networks as local, static objects, to dynamical ones that vary across space and time, has prompted several methodological efforts. First, tools to study spatial, temporal, and spatio-temporal variation of ecological networks in space and in relationship to environmental gradients have been developed and continuously expanded (Poisot *et al.* 2012, 2015, 2017). Second, there has been an improvement in large-scale data-collection, through increased adoption of molecular biology tools (Evans *et al.* 2016, Eitzinger *et al.* 2019, Makiola *et al.* 2019) and crowd-sourcing of data collection (Pocock *et al.* 2015, Bahlai & Landis 2016, Roy *et al.* 2016). Finally, there has been a surge in the development of tools that allow us to *infer* species interactions (Morales-Castilla *et al.* 2015, Dallas *et al.* 2017) based on limited but complementary data on existing network properties (Stock *et al.* 2017), species traits (Gravel *et al.* 2013, Bartomeus *et al.* 2016, Brousseau *et al.* 2017, Desjardins-Proulx *et al.* 2017), and environmental conditions (Gravel *et al.* 2018). These latter approaches tend to perform well in data-poor environments (Beauchesne *et al.* 2016), and can be combined through ensemble modelling or model averaging to generate more robust predictions (Pomeranz *et al.* 2018). The task of inferring interactions is particularly important because ecological networks are difficult to adequately sample in nature (Banašek-Richter *et al.* 2004, Gibson *et al.* 2011, Chacoff *et al.* 2012, Jordano 2016). The common goal to these efforts is to facilitate the prediction of network structure, particularly over space (Poisot & Gravel *et al.* 2016, Gravel *et al.* 2018, Albouy *et al.* 2019) and into the future (Albouy *et al.* 2014), in order to appraise the response of that structure to possible environmental changes.

These disparate methodological efforts share another important trait: their continued success depends on state-of-the art data management, but also on the availability of data that are representative to the area we pretend to model. Novel quantitative tools demand a higher volume of network data; novel collection techniques demand powerful data repositories; novel inference tools demand easier integration between different types of data, including but not limited to: interactions, species traits, taxonomy, occurrences, and local bioclimatic conditions. In short, advancing the science of ecological networks requires us not only to increase the volume of available data, but to pair these data with ecologically relevant metadata. Such data should also be made available in a way that facilitates programmatic interaction so that they can be used by reproducible data analysis pipelines. Poisot & Baiser *et al.* (2016) introduced `mangal.io` as a first step in this direction. In the years since the tool was originally published, we continued development of the data representation, amount and richness of metadata, and digitized and standardized as much ecological data as we could find. The second major release of this database contains over 1300 networks, 120000 interactions across close to 7000 taxa, and represents what is to our best knowledge the most complete collection of species interactions available.

Here we ask if the current mangal database is fit for the purpose of global-scale synthesis research into ecological networks. We conclude that interactions over most of the planet’s surface are poorly described, despite an increasing amount of available data, due to temporal and spatial biases in data collection and digitization. In particular, Africa, South America, and most of Asia have very sparse coverage. This suggests that synthesis efforts on the worldwide structure or properties of ecological networks will be weaker within these areas. To improve this situation, we should digitize available network information and prioritize sampling towards data-poor locations.

## 2 Global trends in ecological networks description

### 2.1 Network coverage is accelerating but spatially biased

The earliest recorded ecological networks date back to the late nineteenth century, with a strong increase in the rate of collection around the 1980s (fig. 1). Although the volume of available networks has increased over time, the sampling of these networks in space has been uneven. In fig. 2, we show that globally, network collection is biased towards the Northern hemisphere, and than different types of interactions have been sampled in different places. As such, it is very difficult to find a spatial area of sufficiently large size in which we have networks of predation, parasitism, and mutualism. The inter-tropical zone is particularly data-poor, either because data producers from the global South correctly perceive massive re-use of their data by Western world scientists as a form of scientific neo-colonialism (as advanced by Mauthner & Parry 2013), thereby providing a powerful incentive *against* their publication, or because ecological networks are subject to the same data deficit that is affecting all fields on ecology in the tropics (Collen *et al.* 2008). As Bruna (2010) identified almost ten years ago, improved data deposition requires an infrastructure to ensure they can be repurposed for future research, which we argue is provided by `mangal.io` for ecological interactions.

**Figure 1:**
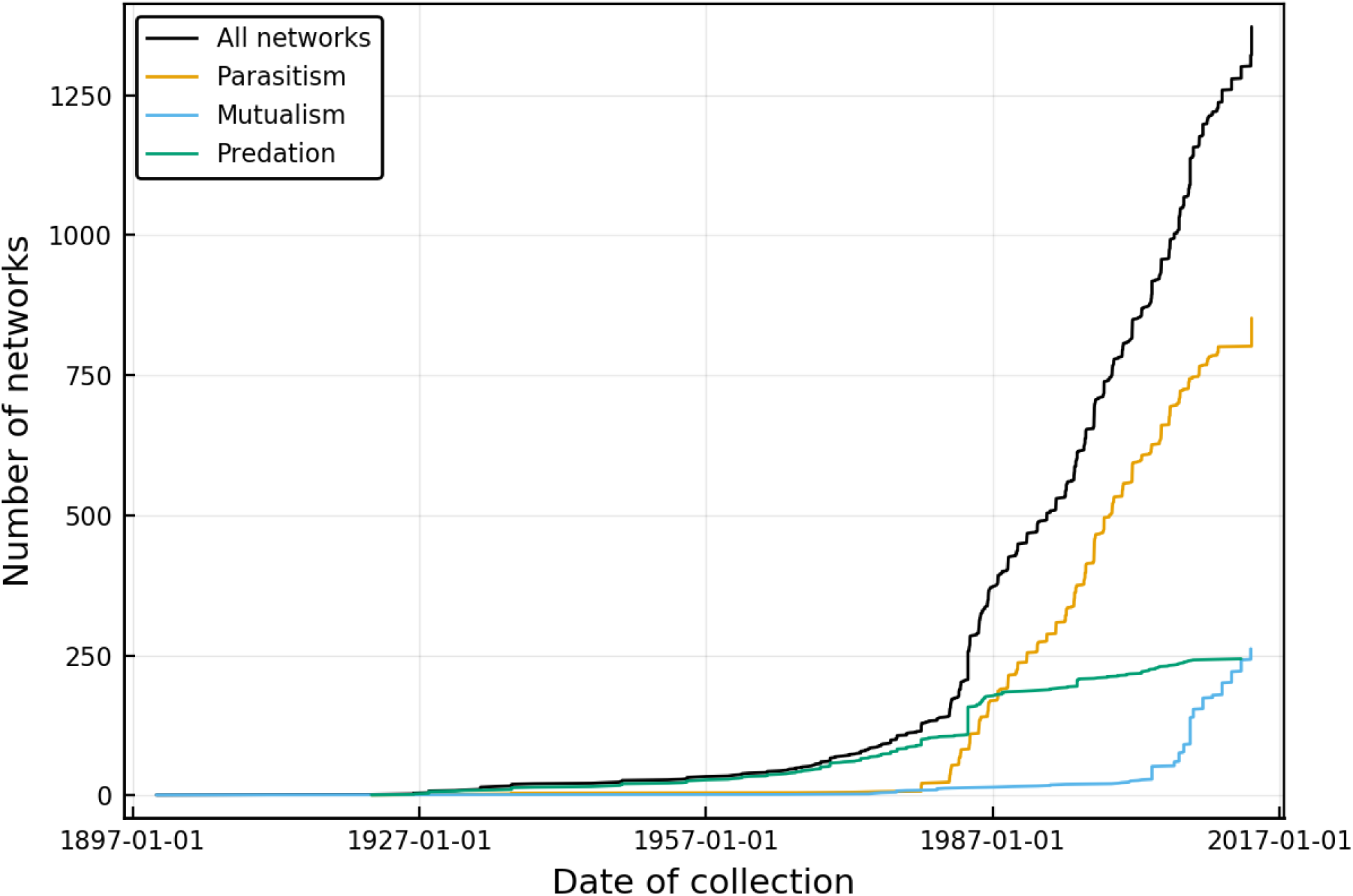
Cumulative number of ecological networks available in `mangal.io` as a function of the date of collection. About 1000 unique networks have been collected between 1987 and 2017, a rate of just over 30 networks a year. This temporal increase proceeds at different rates for diferent types of networks; while the description of food webs is more or less constant, the global acceleration in the dataset is due to increased interest in host-parasite interactions starting in the late 1970s, while mutualistic networks mostly started being recorded in the early 2000s.

**Figure 2:**
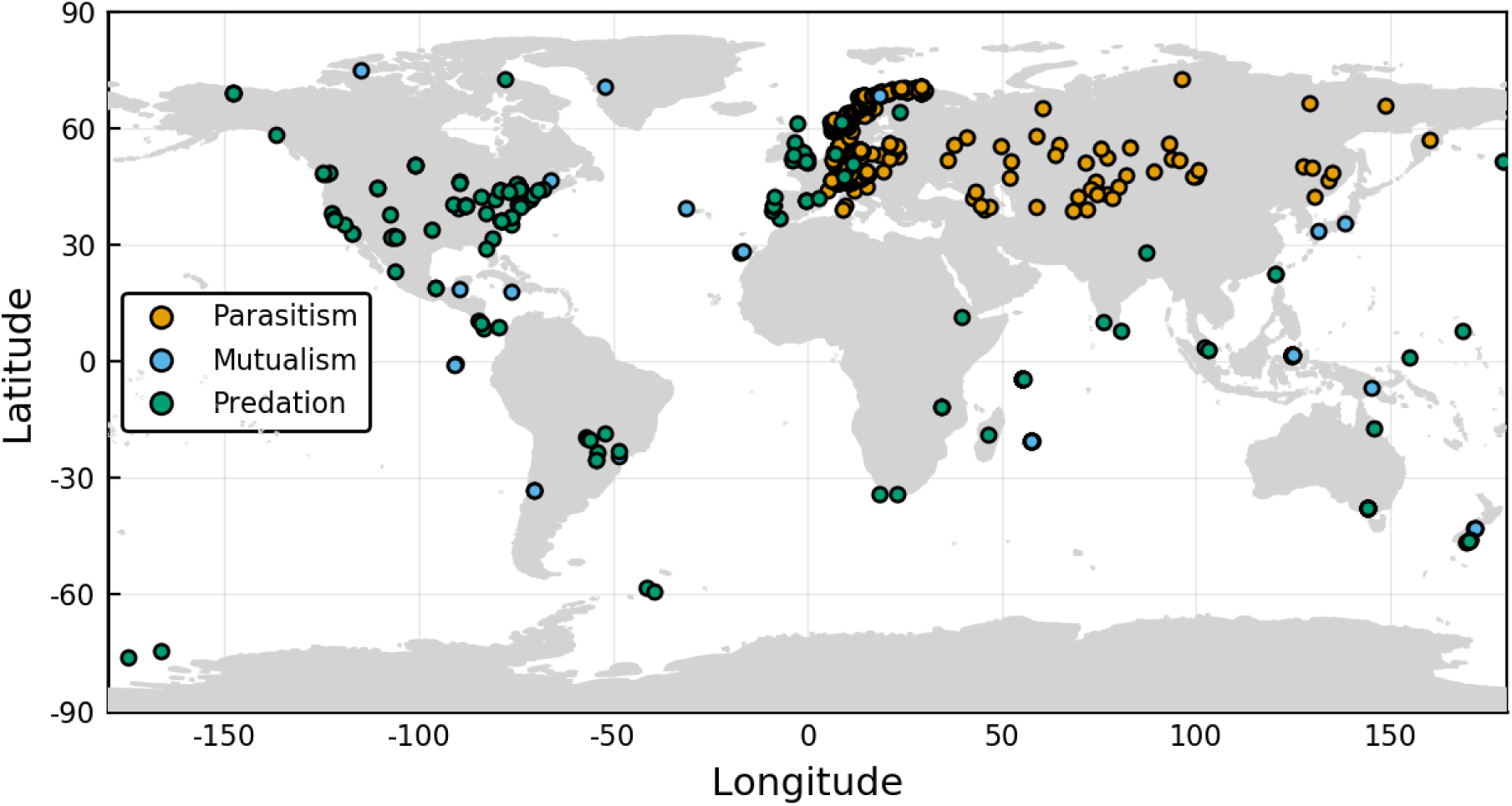
Each point on the map corresponds to a network with parasitic, mutualistic, and predatory interactions. It is noteworthy that the spatial coverage of these types of interactions is uneven; the Americas have almost no parasitic network, for example. Some places have barely been studied at all, including Africa and Eastern Asia. This concentration of networks around rich countries speaks to an inadequate coverage of the diversity of landscapes on Earth.

### 2.2 Different interaction types have been studied in different biomes

Whittaker (1962) suggested that natural communities can be partitioned across biomes, largely defined as a function of their relative precipitation and temperature. For all networks for which the latitude and longitude was known, we extracted the value temperature (BioClim1, yearly average) and precipitation (BioClim12, total annual) from the WorldClim 2 data (Fick & Hijmans 2017). Using these we can plot every network on the map of biomes drawn by Whittaker (1962) (note that because the frontiers between biomes are not based on any empirical or systematic process, they have been omitted from this analysis). In fig. 3, we show that even though networks capture the overall diversity of precipitation and temperature, types of networks have been studied in sub-spaces only. Specifically, parasitism networks have been studied in colder and drier climates; mutualism networks in wetter climates; predation networks display less of a bias.

**Figure 3:**
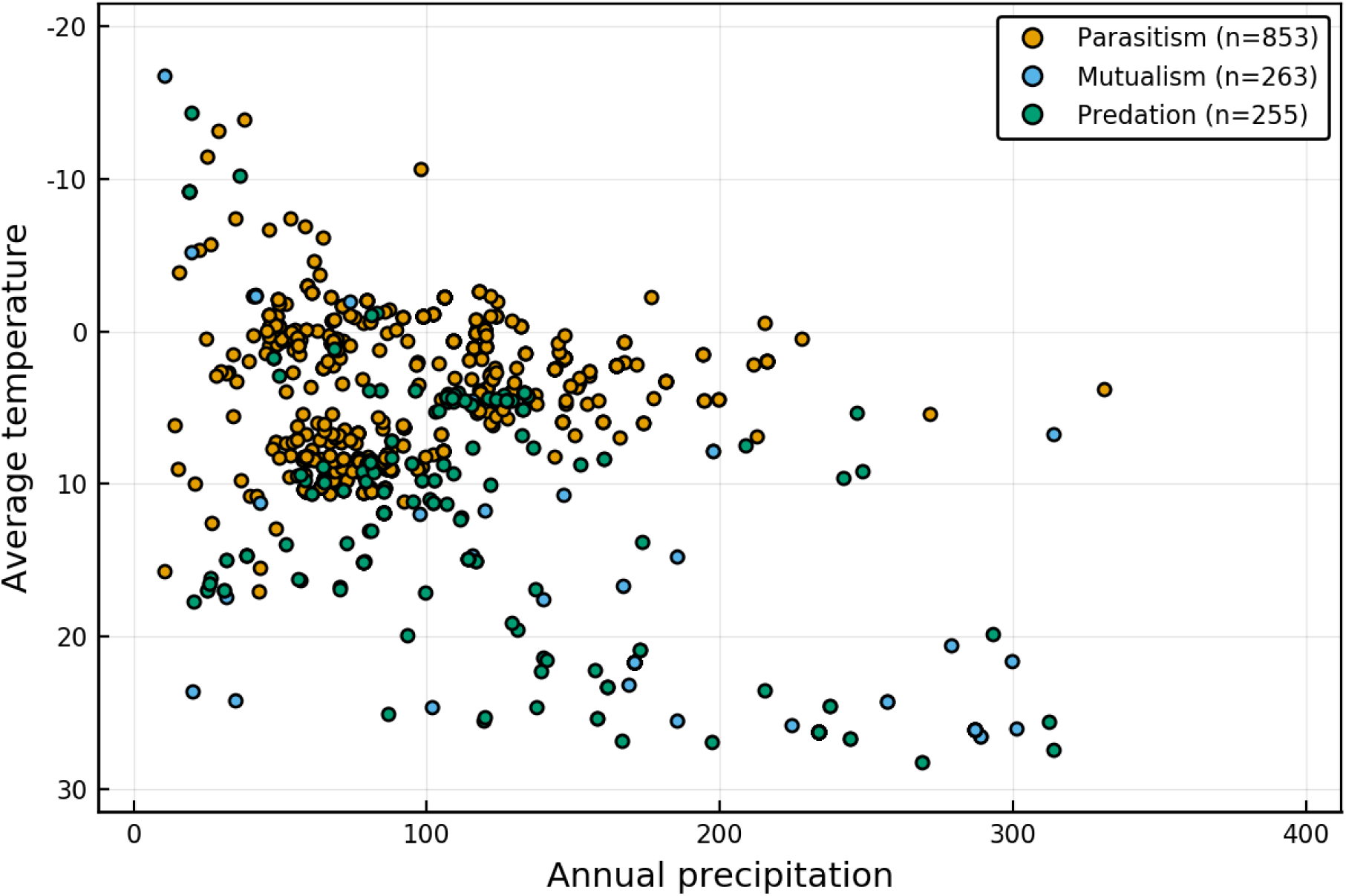
List of networks across in the space of biomes as originally presented by Whittaker (1962). Predation networks, *i.e.* food webs, seem to have the most global coverage; parasitism networks are restricted to low temperature and low precipitation biomes, congruent with the majority of them being in Western Europe.

Scaling up this analysis to the 19 BioClim variables in Fick & Hijmans (2017), we extracted the position of every network in the bioclimatic space, conducted a principal component analysis on the scaled bioclimatic variables, and measured their distance to the centre of this space (**0**). This is a measurement of the “rarity” of the bioclimatic conditions in which any networks were sampled, with larger values indicating more unique combinations (the distance was ranged to [0; 1] for the sake of interpretation). As shown in fig. 4, mutualistic interactions tend to have values that are higher than both parasitism and predation, suggesting that they have been sampled in more unique environments.

**Figure 4:**
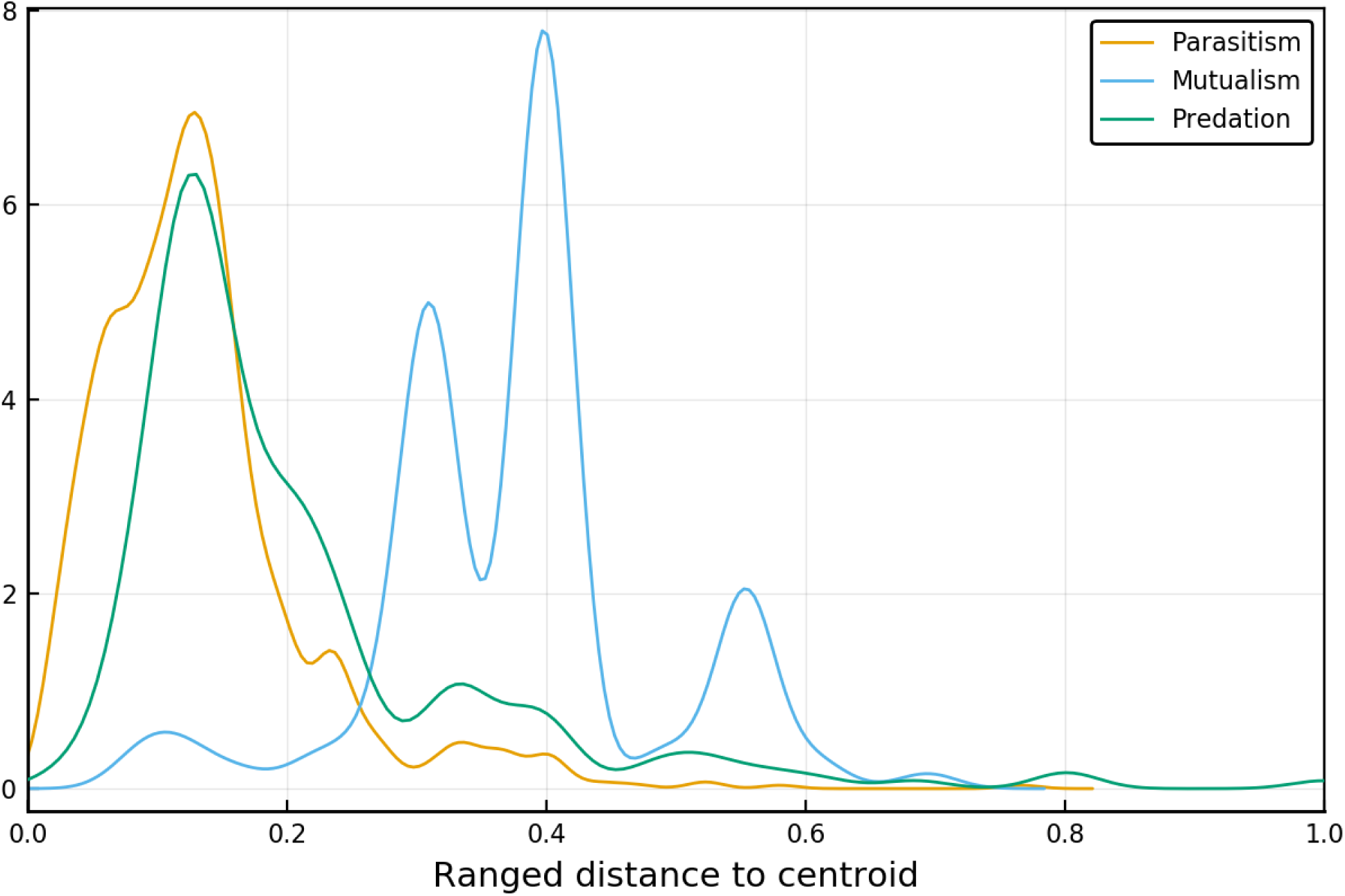
Distance to the centroid (in the scaled climatic space) for each network, as a function of the type of interaction. Larger values indicate that the network is far from its centroid, and therefore represents sampling in a more “unique” location. Mutualistic interactions have been, on average, studied in more diverse locations that parasitism or predatory networks.

### 2.3 Some locations on Earth have no climate analogue

In figures 5, we represent the environmental distance between every pixel covered by *BioClim* data, and the three networks that were sampled in the closest environmental conditions (this amounts to a *k* nearest neighbors with *k* = 3). In short, higher distances correspond to pixels on Earth for which no climate analogue network exists, whereas the darker areas are well described. It should be noted that the three types of interactions studied here (mutualism, parasitism, predation) have regions with no analogues in different locations. In short, it is not that we are systematically excluding some areas, but rather than some type of interactions are more studied in specific environments. This shows how the lack of global coverage identified in fig. 3, for example, can cascade up to the global scale. These maps serve as an interesting measure of the extent to which spatial predictions can be trusted: any extrapolation of network structure in an area devoid of analogues should be taken with much greater caution than an extrapolation in an area with many similar networks.

**Figure 5:**
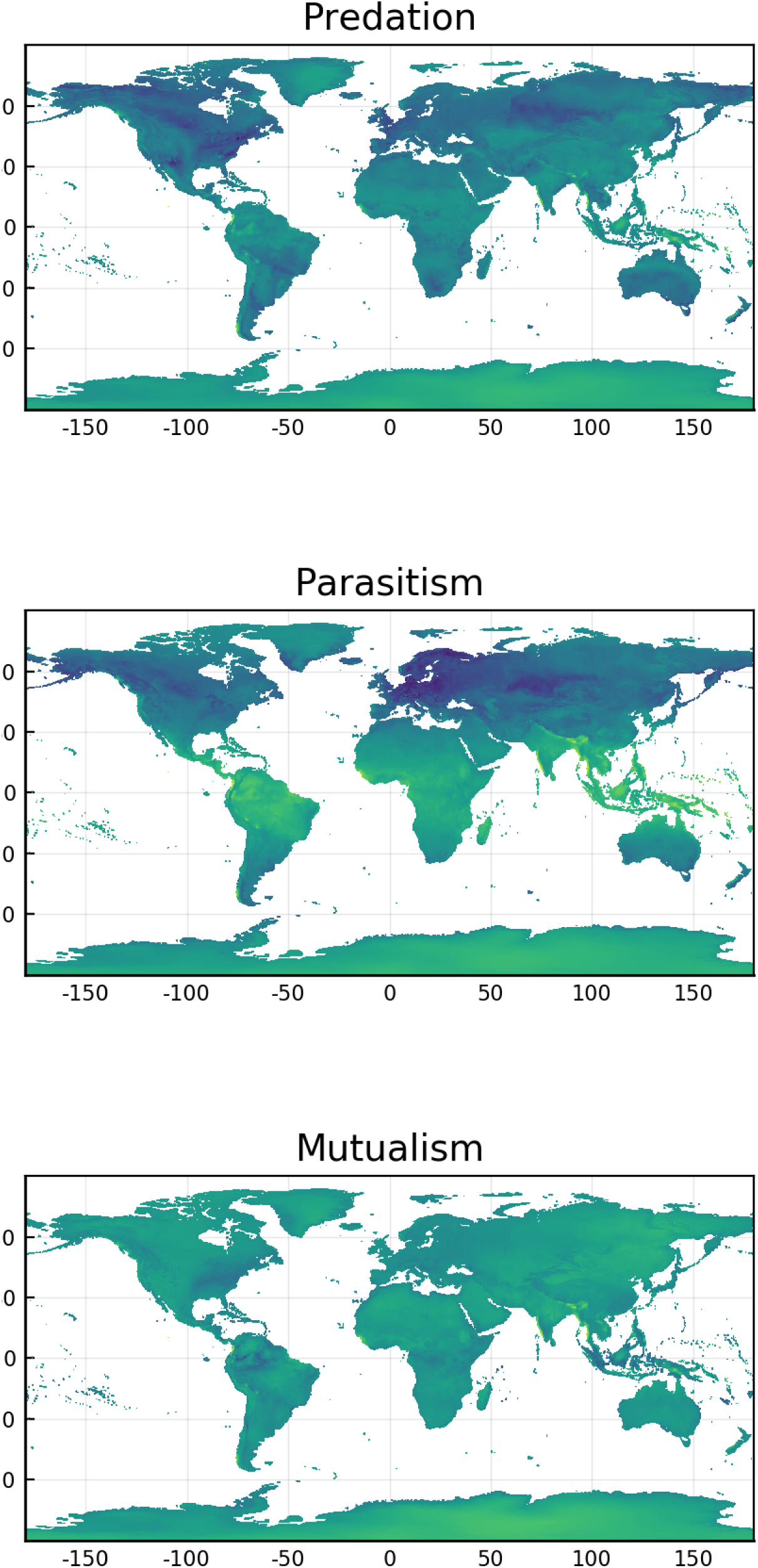
Environmental distance for every terrestrial pixel to its three closest networks. Areas of more yellow coloration are further away from any sampled network, and can therefore not be well predicted based on existing empirical data. Areas with a dark blue coloration have more analogues. The distance is expressed in arbitrary units and is relative.

## 3 Conclusions

### 3.1 For what purpose are global ecological network data fit?

What can we achieve with our current knowledge of ecological networks? The overview presented here shows a large and detailed dataset, compiled from almost every major biome on earth. It also displays our failure as a community to include some of the most threatened and valuable habitats in our work. Gaps in any dataset create uncertainty when making predictions or suggesting causal relationships. This uncertainty must be measured by users of these data, especially when predicting over the “gaps” in space or climate that we have identified. In this paper we are not making any explicit recommendations for synthesis workflows. Rather we argue that this needs to be a collective process, a collaboration between data collectors (who understand the deficiencies of these data) and data analysts (who understand the needs and assumptions of network methods).

One line of research that we feel can confidently be pursued lies in extrapolating the structure of ecological networks over gradients, not at the level of species and their interactions, but at that of the community. Mora *et al.* (2018) revealed that all food webs are more or less built upon the same structural backbone, which is in part due to strong evolutionary constraints on the establishment of species interactions (Dalla Riva & Stouffer 2015); in other words, most networks are expected to be variations on a shared theme, and this facilitates the task of predicting the overarching structure greatly. Finally, this approach to prediction which neglects the composition of networks is justified by the fact that even in the presence of strong compositional turnover, network structure tends to be maintained at very large spatial scales (Dallas & Poisot 2017).

### 3.2 Can we predict the future of ecological networks under climate change?

Perhaps unsurprisingly, most of our knowledge on ecological networks is derived from data that were collected after the 1990s (fig. 1). This means that we have worryingly little information on ecological networks before the acceleration of the climate crisis, and therefore lack a robust baseline. Dalsgaard *et al.* (2013) provide strong evidence that the extant shape of ecological networks emerged in part in response to historical trends in climate change. The lack of reference data before the acceleration of the effects of climate change is of particular concern, as we may be deriving intuitions on ecological network structure and assembly rules from networks that are in the midst of important ecological disturbances. Although there is some research on the response of co-occurrence and indirect interactions to climate change (Araújo *et al.* 2011, Losapio & Schöb 2017), these are a far cry from actual direct interactions; similarly, the data on “paleo-foodwebs”, *i.e.* from deep evolutionary time (Nenzén *et al.* 2014, Yeakel *et al.* 2014, Muscente *et al.* 2018) represent the effect of more progressive change, and may not adequately inform us about the future of ecological networks under severe climate change. However, though we lack baselines against which to measure the present, as a community we are in a position to provide one for the future. Climate change will continue to have important impacts on species distributions and interactions for at least the next century. The Mangal database provides a structure to organize and share network data, creating a baseline for future attempts to monitor and adapt to biodiversity change.

Possibly more concerning is the fact that the spatial distribution of sampled networks shows a clear bias towards the Western world, specifically Western Europe and the Atlantic coasts of the USA and Canada (fig. 2). This problem can be somewhat circumvented by working on networks sampled in places that are close analogues of those without direct information (almost all of Africa, most of South America, a large part of Asia). However, 5 suggests that this approach will rapidly be limited: the diversity of bioclimatic combinations on Earth leaves us with some areas lacking suitable analogues. These regions are expected to bear the worst of the socio-economical (*e.g.* Indonesia) or ecological (*e.g.* polar regions) consequences of climate change. Cameron *et al.* (2019) reached a similar conclusion by focusing on food webs, and our analysis suggests that this worrying trend is in fact one that is shared by almost all types of interactions. All things considered, our current knowledge about the structure of ecological networks at the global scale leaves us under-prepared to predict their response to a warming world. From the limited available evidence, we can assume that ecosystem services supported by species interactions will be disrupted (Giannini *et al.* 2017), in part because the mismatch between interacting species will increase (Damien & Tougeron 2019) alongside the climatic debt accumulated within interactions (Devictor *et al.* 2012).

### 3.3 Active development and data contribution

This is an open-source project: all data and all code supporting this manuscript are available on the Mangal project GitHub organization, and the figures presented in this manuscript are themselves packaged as a self-contained analysis which can be run at any time. Our hope is that the success of this project will encourage similar efforts within other parts of the ecological community. In addition, we hope that this project will encourage the recognition of the contribution that software creators make to ecological research.

One possible avenue for synthesis work, including the contribution of new data to Mangal, is the use of these published data to supplement and extend existing ecological network data. This “semi-private” ecological synthesis could begin with new data collected by authors – for example, a host-parasite network of lake fish in Africa, or a pollination network of hummingbirds in Brazil. Authors could then extend their analyses by including a comparison to analogous data made public in Mangal. After publication of the research paper, the original data could themselves be uploaded to Mangal. This enables the reproducibility of this particular published paper. Even more powerfully, it allows us to build a future of dynamic ecological analyses, wherein analyses are automatically re-done as more data get added. This would allow a sort of continuous assessment of proposed ecological relationships in network structure. This cycle of data discovery and reuse is an example of the Data Life Cycle (Michener 2015) and represents one way to practice ecological synthesis.

Finally, it must be noted that as the amount of empirical evidence grows, so too should our understanding of existing relationships between network properties, networks properties and space, and the interpretation to be drawn from them. In this perspective, the idea of continuously updated analyses is very promising. Following the template laid out by White *et al.* (2019) and Yenni *et al.* (29-Jan-2019), it is feasible to update a series of canonical analyses any time the database grows, in order to produce living, automated synthesis of ecological networks knowledge. To this end, the mangal database has been integrated with `EcologicalNetworks.jl` (Poisot *et al.* 2019), which allows the development of flexible networks analysis pipelines. One immediate target would be to borrow the methodology from Carlson *et al.* (2019), and provide estimate of the sampling effort required to accurately describe combinations of interaction types and bioclimatic conditions.

## Data and code availability

All code is available openly at https://github.com/PoisotLab/MangalSamplingStatus, and the data can be retrieved from `mangal.io` and the BioClim database using the specified files. In addition, weekly updated pages presenting the analyses reported in this manuscript, including the data files, are available at https://poisotlab.github.io/MangalSamplingStatus/.

